# Murine deficiency of peroxisomal L-bifunctional protein (EHHADH) causes medium-chain 3-hydroxydicarboxylic aciduria and perturbs hepatic cholesterol homeostasis

**DOI:** 10.1101/2021.03.02.433634

**Authors:** Pablo Ranea-Robles, Sara Violante, Carmen Argmann, Tetyana Dodatko, Dipankar Bhattacharya, Hongjie Chen, Chunli Yu, Scott L. Friedman, Michelle Puchowicz, Sander M. Houten

**Affiliations:** Department of Genetics and Genomic Sciences, Icahn Institute for Data Science and Genomic Technology, Icahn School of Medicine at Mount Sinai, New York, NY 10029, USA; The Donald B. and Catherine C. Marron Cancer Metabolism Center, Memorial Sloan Kettering Cancer Center, New York, NY 10065, USA; Division of Liver Diseases, Department of Medicine, Icahn School of Medicine at Mount Sinai, New York, NY 10029, USA; Mount Sinai Genomics, Inc, Stamford, CT 06902, USA; Department of Nutrition, School of Medicine, Case Western Reserve University, Cleveland, OH 44106 USA; Department of Pediatrics, University of Tennessee Health Science Center, Memphis, TN 38163, USA

**Keywords:** multifunctional enzyme 1, fatty acid oxidation, omega-oxidation, dodecanedioic acid, hexadecanedioic acid

## Abstract

Peroxisomes play an essential role in the β-oxidation of dicarboxylic acids (DCAs), which are metabolites formed upon ω-oxidation of fatty acids. Genetic evidence linking transporters and enzymes to specific DCA β-oxidation steps is generally lacking. Moreover, the physiological functions of DCA metabolism remain largely unknown. In this study, we aimed to characterize the DCA β-oxidation pathway in human cells, and to evaluate the biological role of DCA metabolism using mice deficient in the peroxisomal L-bifunctional protein (*Ehhadh* KO mice). *In vitro* experiments using HEK-293 KO cell lines demonstrate that ABCD3 and ACOX1 are essential in DCA β-oxidation, whereas both the bifunctional proteins (EHHADH and HSD17B4) and the thiolases (ACAA1 and SCPx) have overlapping functions and their contribution may depend on expression level. We also show that medium-chain 3-hydroxydicarboxylic aciduria is a prominent feature of EHHADH deficiency in mice most notably upon inhibition of mitochondrial fatty acid oxidation. Using stable isotope tracing methodology, we confirmed that products of peroxisomal DCA β-oxidation can be transported to mitochondria for further metabolism. Finally, we show that, in liver, *Ehhadh* KO mice have increased mRNA and protein expression of cholesterol biosynthesis enzymes with decreased (in females) or similar (in males) rate of cholesterol synthesis. We conclude that EHHADH plays an essential role in the metabolism of medium-chain DCAs and postulate that peroxisomal DCA β-oxidation is a regulator of hepatic cholesterol biosynthesis.

## Introduction

Fatty acid β-oxidation (FAO) constitutes the primary source of energy during metabolically demanding conditions such as fasting or exercise (Houten et al. 2016). The major FAO pathway is mitochondrial, but peroxisomes also harbor FAO pathways that metabolize specific carboxylic acids such as very long-chain fatty acids, branched-chain fatty acids, bile acids and long-chain dicarboxylic acids (DCAs) (Wanders & Waterham 2006). These DCAs are products of fatty acid ω-oxidation, an alternative pathway for fatty acid metabolism initiated by microsomal cytochrome P450 enzymes of the CYP4A family (Hardwick et al. 1987; PREISS & BLOCH 1964). CYP4A enzymes catalyze the ω-hydroxylation of long-chain fatty acids that are subsequently converted into long-chain DCAs by the successive action of an alcohol and an aldehyde dehydrogenase. Peroxisomal β-oxidation of DCAs generates unique products such as adipic and succinic acid (Jin et al. 2015; Rusoff et al. 1960; Tserng & Jin 1991a), the latter being able to enter the TCA cycle. In addition, peroxisomal β-oxidation yields acetyl-CoA units, which have a variety of fates including lipogenesis, ketogenesis, or oxidation in the TCA cycle. These metabolic pathway connections suggest an important role for DCA β-oxidation in intermediary metabolism.

Mitochondrial and peroxisomal FAO as well as ω-oxidation are usually induced in parallel, particularly in the liver and the kidney, under conditions of high lipid flux such as during fasting (Mortensen & Gregersen 1981; Pettersen et al. 1972). Notably, medium-chain dicarboxylic aciduria; i.e. the excretion of C6-, C8-, and C10-DCAs (adipic, suberic and sebacic acids, respectively) in the urine, is a feature shared between fasting individuals (ketosis) and patients with inherited defects in mitochondrial FAO (Fontaine et al. 1998; Gregersen et al. 1980; Karpati et al. 1975; Przyrembel et al. 1976; Röschinger et al. 2000; Vianey-Saban et al. 1998). These findings further illustrate the important role of fatty acid ω-oxidation and subsequent DCA β-oxidation under lipid metabolic stress. However, the physiological roles of DCA metabolism remain largely to be established.

DCA metabolism begins with their activation by a dicarboxylyl-CoA synthetase (Vamecq et al. 1985). Subsequent DCA β-oxidation is thought to occur in peroxisomes (Dirkx et al. 2007; Ferdinandusse et al. 2004; Kølvraa & Gregersen 1986; Suzuki et al. 1989). Both EHHADH (also known as peroxisomal L-bifunctional protein, LBP) and HSD17B4 (also known as D-bifunctional protein, DBP) can catalyze the second and third steps of peroxisomal FAO. We and others have demonstrated that EHHADH plays a prominent role in the oxidation of long-chain DCAs (Ding et al. 2013; Dirkx et al. 2007; Ferdinandusse et al. 2004; Houten et al. 2012; Nguyen et al. 2008). Other peroxisomal proteins that participate in DCA metabolism are the ABCD3 transporter (van Roermund et al. 2014), ACOX1 (acyl-CoA oxidase 1) (Ferdinandusse et al. 2004; Poosch & Yamazaki 1989), and the thiolases ACAA1 (3-ketoacyl-CoA thiolase) and SCPx (sterol carrier protein X) (Ferdinandusse et al. 2004). In many cases, genetic evidence that links the encoding genes to DCA β-oxidation is lacking. In summary, ω-oxidation and subsequent peroxisomal DCA β-oxidation is an alternate pathway for fatty acid metabolism that does not rely on the classic mitochondrial FAO pathway.

In this study, we assessed the contribution of candidate peroxisomal proteins to DCA β-oxidation using KO HEK-293 cell lines created by CRISPR-Cas9 genome editing. We also examined the physiological role of DCA β-oxidation through the study of the *Ehhadh* KO mouse model. Finally, we assessed the relevance of peroxisomal DCA metabolism in mouse models with mitochondrial FAO defects. The results of our study further highlight an important role for peroxisomal DCA β-oxidation in intermediary metabolism.

## Material and Methods

### Materials

HEK-293 cells (ATCC^®^ CRL-1573™) were obtained from the American Type Culture Collection (Manassas, VA, USA). Dulbecco’s modified Eagle’s medium (DMEM), Krebs–Henseleit buffer, William’s E Medium with GlutaMAX™ Supplement (WEGG), heat-inactivated fetal bovine serum (FBS), penicillin, streptomycin, and Lipofectamine 2000 were purchased from Thermo Fisher Scientific (Waltham, MA, USA). CRISPR-Cas9 plasmid [pSpCas9(BB)-2A–green fluorescent protein (GFP); PX458] (Ran et al. 2013) was obtained from Addgene (Watertown, MA, USA). The blocking buffer and the fluorescent secondary anti-rabbit and anti-mouse IgG antibodies were obtained from LI-COR Biosciences (Lincoln, NE, USA). [^2^H]-water (99% enriched), the L-carnitine internal standard (^2^H_9_-carnitine) and the acylcarnitine internal standard mix containing ^2^H_9_-carnitine, ^2^H_3_-acetylcarnitine (C2), ^2^H_3_-propionylcarnitine (C3), ^2^H_3_-butyrylcarnitine (C4), ^2^H_9_-isovalerylcarnitine (C5), ^2^H_3_-octanoylcarnitine (C8), ^2^H_9_-myristoylcarnitine (C14), and ^2^H_3_-palmitoylcarnitine (C16), were from Cambridge Isotope Laboratories (Tewksbury, MA, USA). Minimal essential medium, bovine serum albumin (fatty acid free), L-carnitine, lauric acid (C12:0), [U-^13^C]-dodecanedioic acid ([U-^13^C]-C12-DCA, 99 atom % ^13^C), dodecanedioic acid (C12-DCA), and hexadecanedioic acid (C16-DCA) were obtained from MilliporeSigma (Burlington, MA, USA). L-Aminocarnitine [L-AC; (R)-aminocarnitine, minimum 97%], 3-OH-adipic (3-hydroxyhexanedioic acid, 97%), 3-OH-suberic (3-hydroxyoctanedioic acid, 97%), and 3-hydroxydodecanedioic (97%) were obtained from Toronto Research Chemical Inc. (Toronto, ON, Canada). 3-OH-sebacic (>98%) was obtained from Santa Cruz (Dallas, TX). All other chemicals were of analytical grade.

### Animals

All mouse experiments were approved by the Institutional Animal Care and Use Committee (IACUC) of the Icahn School of Medicine at Mount Sinai (Protocols: IACUC-2014-0100 and IACUC-2014-0061) and comply with the Guide for the Care and use of Laboratory Animals (NIH Publications No. 8023, 8^th^ edition, 2011). Mice were housed on a 12h light/dark cycle. Mice were fed with regular chow (PicoLab Rodent Diet 20; LabDiet, St. Louis, MO, USA) and studied in the adult stage. *Ehhadh* KO mice were provided by Dr. Janardan K. Reddy (Qi et al. 1999) and backcrossed to a C57BL/6N background (Violante et al. 2019). The generation and acquisition of the LCAD KO (*Acadl* KO) mouse has been described (Houten et al. 2013; Kurtz et al. 1998). The generation of *Ehhadh/Acadl* double KO mice (DKO) is described in the Results section. The *Acadl* allele was genotyped using an SNP-based genotyping assay (Luther et al 2012) and the *Ehhadh* allele was genotyped using a PCR assay.

We used three experimental cohorts. For the DKO experiments, the cohort consisted of 32 WT (16 males, 16 females, average age = 5.4 ± 1.7 months), 33 *Ehhadh* KO (12 males, 21 females, average age = 5.6 ± 2.5 months), 12 *Acadl* KO (2 males, 10 females, average age = 10.3 ± 1.7 months), and 6 DKO mice (3 males, 3 females, average age = 4.3 ± 0.8 months). For the cholesterol synthesis experiment, the cohort consisted of 11 WT (5 males, 6 females, average age = 3.5 ± 1.0 months) and 12 *Ehhadh* KO (6 males, 6 females, average age = 3.1 ± 1.3 months) mice. The cohort used for the urinary DCA data with or without L-AC consisted of 8 WT (1 male, 7 females, average age = 6.1 ± 0.4 months) and 10 *Ehhadh* KO (2 males, 8 females, average age = 5.7 ± 0.9 months) mice that were fasted for 8 hours. We also used unpublished plasma acylcarnitine data of a previously reported L-AC cohort that included WT and *Ehhadh* KO mice subjected to overnight food withdrawal (Violante et al. 2019).

Overnight food withdrawal was performed by removing the food and placing the mice in a clean cage at 6 p.m. just before the day/night light cycle switches. Early the next morning, blood glucose was measured using Bayer Contour blood glucose strips. Spontaneously voided urine was collected by scruffing the mice and placing them over a plastic wrap in order to collect the specimen. Mice were euthanized by exposure to CO_2_. Immediately afterwards, blood was collected for the preparation of EDTA plasma. Organs were snap-frozen in liquid nitrogen and stored at −80°C for future analysis. Plasma was used for acylcarnitine analysis. Urine was used for organic acid analysis to measure DCA content.

For cholesterol synthesis, animals were injected with deuterium oxide (^2^H_2_O) as 0.9% NaCl at 7 a.m., aiming to reach 4% enrichment of total body water using the following formula: *Volume of heavy water injected (mL) = Body weight (g) * 0.75 (Water content in the body) * 0.04 (% of enrichment)*. After 7 hours, a small amount of blood was drawn from the animals to measure ^2^H_2_O enrichment, and immediately after, the mice were euthanized by exposure to CO_2_. The liver was quickly frozen with a freeze-clamp, wrapped in aluminum foil, temporarily stored in liquid nitrogen, and ultimately transferred and stored at −80°C.

For the L-AC experiments, animals received an intraperitoneal injection of vehicle (0.9% NaCl) or L-AC (16 mg/kg) at 6 p.m., before the 12-hr dark cycle, followed by overnight food withdrawal. In a second experiment, animals received either vehicle or L-AC (16 mg/kg i.p.) at 9 a.m., followed by food withdrawal for 8 hours. Urine, blood and organs were collected as described above.

### Cell culture and incubation with fatty acids

HEK-293 cells were cultured in DMEM supplemented with 10% FBS, 100 U/mL penicillin, and 100 mg/mL streptomycin, in a humidified atmosphere of 5% CO_2_, at 37°C. The L-AC treatment and subsequent acylcarnitine profiling after lauric acid loading in HEK-293 cells were performed as previously described (Chegary et al. 2008; Violante et al. 2013, 2019). For the measurement of DCA metabolism, HEK-293 cells were seeded in 6-well plates (0.8 × 10^6^ cells per well) and incubated overnight at 37°C. The following day, the medium was removed and 2 mL of the incubation mixture were added to each well. Incubation mixtures consisted of DMEM (with 10% FBS and antibiotics) supplemented with 250 μM hexadecanedioic acid (C16-DCA, added from a 25 mM stock in 100% ethanol) or vehicle. After incubation for 72 hours in a humidified CO_2_ incubator (5% CO_2_, 95% air) at 37°C, the medium was collected and cells were washed and resuspended in 100 μL of RIPA buffer. Protein content was measured using the bicinchoninic acid (BCA) assay and human serum albumin as standard.

### EHHADH overexpression

Control and *HSD17B4* KO cell lines were transfected with a plasmid containing the human *EHHADH* cDNA (MGC premier cDNA clone for EHHADH, pCS6 (BC038948), Transomic) or an empty vector. Transfection was performed with Lipofectamine 2000 following the supplier’s instructions. After 24-hr incubation, cells were incubated with lauric acid +/− L-AC in one experiment, and C16-DCA in the other experiment. The medium was analyzed after 72-h incubation as previously described. Overexpression of EHHADH was confirmed by immunoblot.

### Acylcarnitine profiling

HEK-293 cells, liver slices culture medium, and mouse plasma acylcarnitines were measured after derivatization to butylesters according to a standardized protocol (Violante et al. 2019). Tissue acylcarnitines were measured in freeze-dried tissue samples (approx. 50 mg of liver) after derivatization to propylesters essentially as described (Ranea-Robles et al. 2020; van Vlies et al. 2005).

### Organic acid analysis

Organic acids from mouse urine, HEK-293 cell culture medium, and mouse liver slices culture medium were analyzed essentially as previously described with minor modifications (Leandro et al. 2020). Urine specimens were normalized to creatinine (creatinine range between 0.1 and 1 μmol). Organic acids were oximated prior to acidification and extraction. Quantification of 3-hydroxy DCAs was performed using a calibration curve of a mix containing 3-OH-adipic, 3-OH-suberic, 3-OH-sebacic and 3-OH-dodecanedioic acid. 3-OH-adipic acid was quantified as the sum of the open chain form and the cyclic lactone form (3-hydroxyadipic acid 3,6-lactone).

### Generation of CRISPR-Cas9 KO cell lines

The generation of gene KO HEK-293 cell lines using the CRISPR-Cas9 genome editing technique was performed as previously described (Violante et al. 2019). For each gene, 2 guides were designed (Table S1). We selected 2 independent clonal KO cell lines for each targeted gene (Violante et al. 2019). A list of the newly established cell lines for this study is provided in Table S1.

### Immunoblot analysis

Cells and tissues were lysed in RIPA buffer supplemented with protease and phosphatase inhibitors (Thermo Fisher Scientific), followed by sonication and centrifugation (10 min at 12,000 rpm at 4°C). Protein concentration was determined by the BCA method. Proteins were separated on Bolt™ 4-12% Bis-Tris Plus gels (Invitrogen, Thermo Fisher Scientific), blotted onto a nitrocellulose membrane (926-31092, LI-COR) and detected using the following primary antibodies: anti-ACOX1 (ab184032, Abcam), anti-ACAA1 (12319-2-AP, Proteintech), anti-EHHADH (GTX81126, Genetex), anti-HSD17B4 (ab128565, Abcam), anti-HMGCR (AMAb90618, Atlas Antibodies), anti-citrate synthase (GTX628143, Genetex), and anti-α-tubulin (32-2500, Thermo Fisher). Anti-MVK and anti-PMVK antibodies were a gift from Dr. Hans Waterham (AMC, Amsterdam) (Hogenboom et al. 2004b, a). Proteins were visualized with goat anti-mouse and goat anti-rabbit secondary antibodies IRDye 800CW and IRDye 680RD (LI-COR, 926-32 210, 926-68 070, 926-32 211, 926-68 071). Band intensities were quantified using the Fiji distribution of ImageJ 1.x (Schindelin et al. 2012).

### Measurement of cholesterol synthesis and *de novo* lipogenesis by the ^2^H_2_O method

The contribution of cholesterol synthesis (^2^H-labeled cholesterol) to the pool of total hepatic cholesterol and the contribution of *de novo* lipogenesis (^2^H-labeled TG-bound palmitate, oleate and stearate) to the pool of total hepatic TG-bound fatty acids in mice labeled with ^2^H_2_O were determined using gas chromatography-mass spectrometry (GC-MS), as previously described (Bederman et al. 2009; Brunengraber et al. 2003). These measurements were performed at the Metabolic Phenotyping Mass Spectrometry Core (MPMS), UTHSC, Memphis, TN.

### Metabolic tracing of [U-^13^C]-C12-DCA in precision-cut mouse liver slices

Mouse livers were isolated from female fed WT and *Ehhadh* KO adult mice (n = 5) between 7 and 11 a.m. Precision-cut liver slices (diameter 8 mm, thickness 250 μm) were prepared in ice-cold Krebs-Henseleit buffer saturated with carbogen (95% O_2_ / 5% CO_2_), using the Leica VT1200 S vibrating blade microtome with Vibrocheck as previously described (De Graaf et al. 2010; Zimmermann et al. 2009). After slicing, samples were preincubated 1 hour in WEGG medium, and randomly selected for the vehicle or [U-^13^C]-C12-DCA treatment. Slices were then incubated 24 hours in cell culture inserts in 6-well plates with fresh WEGG medium containing 0.4 mM carnitine and either 250 μM [U-^13^C]-C12-DCA or vehicle. [U-^13^C]-disodium dodecanedioate was prepared by dissolving the acid in ethanol followed by the addition of 1M NaOH in 80% methanol (2 times the equimolar amount). After evaporation of the solvent, the DCA tracer was dissolved in the WEGG medium. Liver slices were stored at −80°C and the medium was stored at −20°C for metabolite analysis.

^13^C-labeling patterns of C6-DCA, C8-DCA, 3-OH-C8-DCA, C6DC-carnitine, C8DC-carnitine, citric acid cycle intermediates, amino acids, and the ketone body β-hydroxybutyrate (β-OHB) were analyzed from the culture medium and liver slice homogenates using stable isotopomer mass spectrometry analysis, as previously described (Zhang et al. 2015). Concentrations of universally ^13^C labeled DCAs and 3-hydroxy DCAs (M+6 in the case of C6-DCAs, and M+8 in the case of C8-DCAs) in culture medium were quantified using the same calibration curves used for the quantification of the correspondent non-labeled metabolites.

The fractional isotopic enrichments (via ^13^C label incorporation) for citric acid cycle intermediates and β-OHB were defined as molar percent enrichment (MPE) after background correction using a matrix method, as previously described (Zhang et al. 2015). Briefly, the MPE (unit: percent) is calculated as:

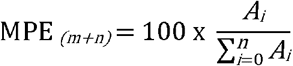

where MPE_(*M+n*)_ is the *n*th ^13^C-labeled mass isotopomer enrichment of each metabolite. *A* is the abundance (arbitrary unit). The variable *i* varied from zero to *n*, while *n* is no more than 6. *A*_(0)_ is the abundance for the non-labeled metabolite, and *A_(i)_* is the abundance for mass isotopomer labeled with *i* ^13^C, regardless of position (Zhang et al. 2015).

### RNA-seq, differential gene expression, pathway enrichment

RNA was isolated using QIAzol lysis reagent, followed by purification using the RNeasy kit (Qiagen). RNA samples were submitted to the Genomics Core Facility at the Icahn Institute and Department of Genetics and Genomic Sciences, and sequenced essentially as previously described (Argmann et al. 2017). Briefly, mRNA-focused cDNA libraries were generated using Illumina reagents (polyA capture), and samples were run on an Illumina HiSeq 2500 sequencer to yield a read depth of approximately 43 million 100 nucleotide single-end reads per sample. Reads from fastq files were aligned to the mouse genome mm10 (GRCm38.75) with STAR (release 2.3.0e) (Dobin et al. 2013) and summarized to gene- and exon-level counts using featureCounts (Liao et al. 2014). Differential gene expression analysis was conducted with R packages limma (Law et al. 2014; Ritchie et al. 2015) and DESeq2 (Love et al. 2014) as previously described (Argmann et al. 2017). The cut-off for differential gene expression was chosen at an adjusted p-value (Benjamini-Hochberg) of < 0.05. Gene Set Enrichment Analysis (GSEA) using the Molecular Signatures Database (MSigDB) was performed to determine whether or not a specific subset of genes were enriched in *Ehhadh* KO livers (Subramanian et al. 2005).

### Statistical analysis

Data are displayed as the mean ± the standard deviation (SD) or the standard error of the mean (SEM) as indicated in the figure legends. In most cases, individual values are shown. Differences were evaluated using unpaired t-test with Welch’s correction, one-way ANOVA with Tukey’s multiple comparison test, or two-way ANOVA, as indicated in the figure legends. If one or more groups were not distributed normally, non-parametric Kruskal-Wallis test with Dunn’s multiple comparison posthoc test was used. Significance is indicated in the figures. GraphPad Prism 8 was used to compute statistical values.

## Results

### Generation and validation of HEK-293 cell lines with peroxisomal enzyme deficiencies

We previously used CRISPR-Cas9 genome editing in HEK-293 cells to demonstrate that peroxisomes can oxidize medium- and long-chain fatty acids through a pathway involving ABCD3 and HSD17B4 (Violante et al. 2019). In our assay, carnitine palmitoyltransferase 2 (CPT2) inhibition using LAC in combination with lauric acid (C12) loading is used to unveil the production of peroxisomal acylcarnitines including C10-carnitine. A defect in peroxisomal biogenesis (*PEX13* KO) or peroxisomal fatty acid transport (*ABCD3* KO) abolishes the production of the peroxisomal acylcarnitines (Violante et al. 2019). In a set of newly generated KO cell lines (Table S1), we now study each step in peroxisomal FAO in more detail. We demonstrate that ACOX1 is essential for the production of C10-carnitine, and thus catalyzes the first step of the peroxisomal oxidation of C12-CoA (Fig. 1A). For the fourth and last step of the peroxisomal oxidation of lauric acid, both ACAA1 and SCPx (encoded by the *SCP2* gene) appear non-essential as the single KO cell lines had no defect in C10-carnitine production (Fig. 1A). *ACAA1 /SCP2* DKO cell lines, however, failed to accumulate C10-carnitine showing that these enzymes are redundant in the peroxisomal oxidation of lauric acid (Fig. 1A).

**Fig. 1.**
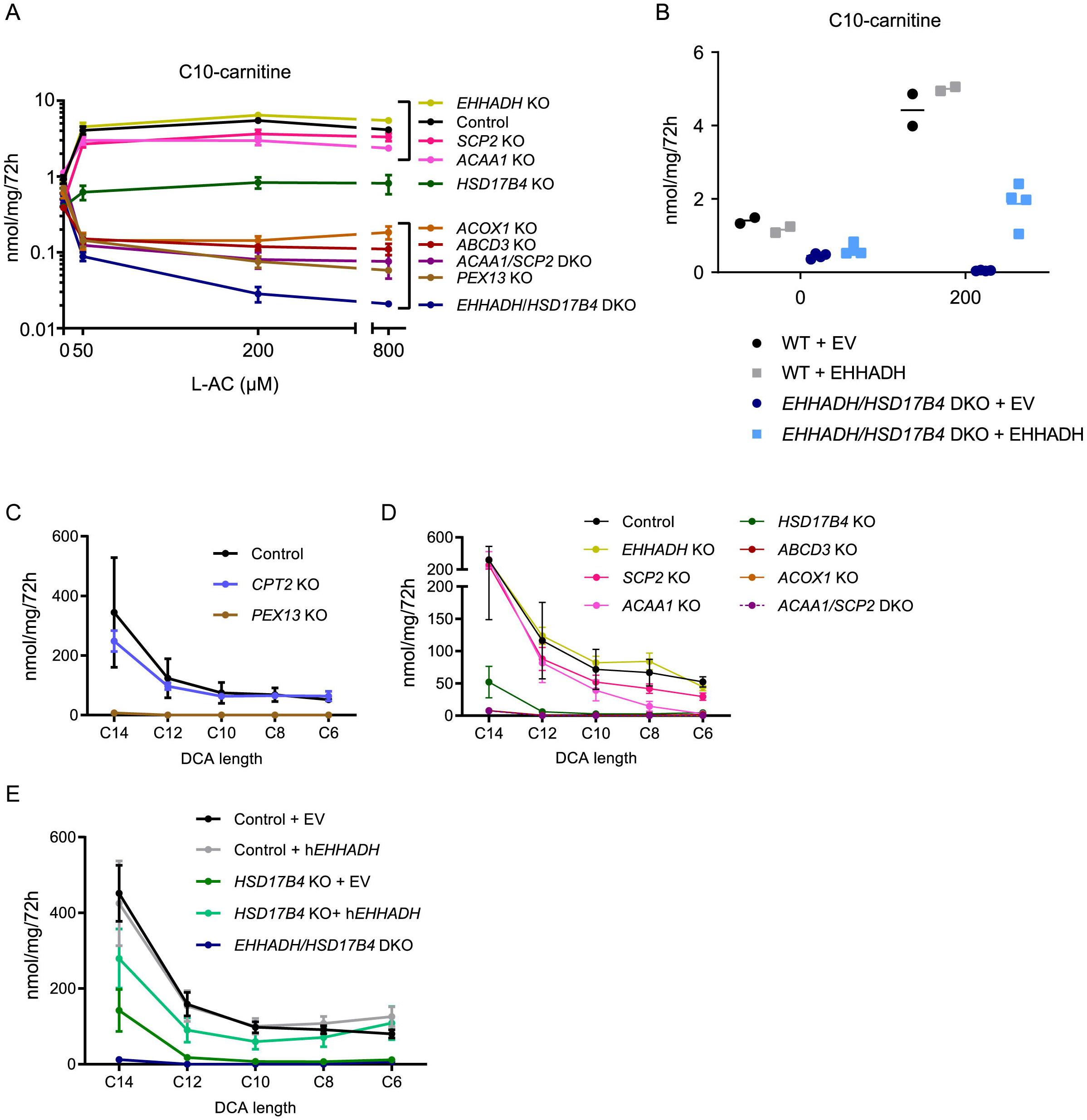
Peroxisomal β-oxidation of lauric acid and hexadecanedioic acid in HEK-293 cells. **A)** Production of peroxisomal C10-carnitine (nmol/mg protein/72h) in the extracellular media of KO cell lines after loading with C12:0 in the presence of increasing concentrations of the CPT2 inhibitor L-AC. The cell lines are placed in 3 groups; those with no apparent defect (top), those with deficient but detectable β-oxidation (only *HSD17B4* KO), and finally, those with no detectable peroxisomal β-oxidation (bottom). Note that the currently reported values are 25 times larger than those reported by Violante et al (Violante et al. 2019). The current values reflect the total production in 500μL media and not the 20μl specimen used for the MS analysis. **B)** Rescue of the production of peroxisomal C10-carnitine (nmol/mg protein/72h) in *EHHADH/HSD17B4* double KO (DKO) cell lines by EHHADH overexpression. **C)** Production of chain-shortened DCAs (C14 to C6, nmol/mg protein/72h) by control, *CPT2* KO and *PEX13* KO HEK-293 cell lines after 72-hr incubation with hexadecanedioic acid (C16-DCA). **D)** Production of chain-shortened DCAs (C14 to C6, nmol/mg protein/72h) by control, *ABCD3* KO, *ACOX1* KO, *EHHADH* KO, *HSD17B4* KO, *ACAA1* KO, *SCP2* KO, and *ACAA1/SCP2* DKO HEK-293 cell lines after 72-hr incubation with C16-DCA. Note that lines for *ABCD3* KO, *ACOX1* KO, and *ACAA1/SCP2* DKO almost completely overlap. **E)** Production of chain-shortened DCAs (C14 to C6, nmol/mg protein/72h) by control, *HSD17B4* KO and *HSD17B4/EHHADH* DKO HEK-293 cell lines after 72-hr incubation with C16-DCA. Control and *HSD17B4* KO cell lines were analyzed with or without overexpression of EHHADH. Data are displayed as the mean ± SEM (**A**), individually + the mean (**B**), or as the mean ± S.D (**C-E**) of 2-4 independent experiments. All experiments were carried out with 2 clonal cell lines.

We also studied in more detail the role of EHHADH and HSD17B4. We previously demonstrated that HSD17B4, but not EHHADH, is essential in the peroxisomal β-oxidation of lauric acid in HEK-293 cells (Violante et al. 2019). The *HSD17B4* KO cell lines, however, produced small amounts of C10-carnitine (Fig. 1A). Therefore, we hypothesized that despite its low expression in HEK-293 cells, EHHADH can partially compensate for the loss of HSD17B4 (Violante et al. 2019). To test this hypothesis, we generated *EHHADH/HSD17B4* DKO cell lines. Production of the peroxisomal C10-carnitine was fully defective in *EHHADH/HSD17B4* DKO cell lines (Fig. 1A), mirroring the profile of *PEX13* KO, *ABCD3* KO, *ACOX1* KO and *ACAA1/SCP2* DKO cell lines. This result indicates that, depending on its expression level, EHHADH may be able to take over the function of HSD17B4 in lauric acid oxidation. To further demonstrate this concept, we overexpressed EHHADH in *EHHADH/HSD17B4* DKO cell lines. EHHADH overexpression rescued defective peroxisomal C10-carnitine production upon L-AC treatment (Fig. 1B). Combined, these results indicate that EHHADH and HSD17B4 are functionally redundant in lauric acid oxidation, and their individual roles may in part be determined by tissue expression levels.

### Hexadecanedioic acid is β-oxidized in the peroxisome of HEK-293 cells

Next, we explored if HEK-293 cells can be used to study the metabolism of DCAs. The generation of DCAs depends on the expression of members of the cytochrome P450 4A family, which is restricted to liver and kidney (Hardwick et al. 1987; Mortensen 1992; Uhlén et al. 2015). Accordingly, expression of *CYP4A11* and *CYP4A22* is very low in HEK-293 cells (Uhlén et al. 2015). We therefore supplied HEK-293 cells directly with hexadecanedioic acid (C16-DCA) and dodecanedioic acid (C12-DCA). For both DCAs we observed production of chain-shortened DCAs in the cell culture media demonstrating that DCA β-oxidation occurs in HEK-293 cells (Fig. 1C, D and data not shown). We next loaded all our mutant cell lines with C16-DCA and determined DCA chain shortening through β-oxidation. DCA chain shortening was defective in *PEX13* KO cell lines, but not affected in *CPT2* KO cell lines (Fig. 1C). This observation is consistent with the notion that DCA β-oxidation occurs primarily in peroxisomes and not in mitochondria.

We then studied the contribution of candidate peroxisomal transporters and enzymes to DCA β-oxidation. We observed deficient DCA β-oxidation in *ABCD3* KO and *ACOX1* KO cell lines (Fig. 1D). Production of chain-shortened DCAs was also severely reduced in *HSD17B4* KO cell lines, whereas it was not affected in *EHHADH* KO cell lines (Fig. 1D). This is surprising as all currently available evidence clearly implicates EHHADH in DCA metabolism (Ding et al. 2013; Dirkx et al. 2007; Ferdinandusse et al. 2004; Houten et al. 2012; Nguyen et al. 2008). As discussed above for the oxidation of lauric acid (Fig. 1A, B), this finding may be related to the low expression level of *EHHADH* in HEK-293 cells (Violante et al. 2019). Notably, some C14-DCA remained detectable in *HSD17B4* KO cell lines indicating a potential small contribution of EHHADH (Fig. 1D). Indeed, *EHHADH/HSD17B4* DKO cell lines were completely defective in C16-DCA β-oxidation (Fig. 1E). To overcome the limitation of low *EHHADH* expression in HEK-293 cells, we overexpressed EHHADH in *HSD17B4* KO cells and evaluated C16-DCA β-oxidation. EHHADH overexpression rescued the defective C16-DCA β-oxidation in *HSD17B4* KO cells (Fig. 1E, Supplementary Fig. S1B). Moreover, EHHADH overexpression slightly increased C6-DCA production in control HEK-293 cells (Fig. 1E). Combined, these data illustrate that both EHHADH and HSD17B4 are able to handle DCAs and that their role may be mainly driven by tissue expression levels.

Production of C14-, C12- and C10-DCA was not affected in *ACAA1* KO cell lines, but the levels of C8- and C6-DCA were clearly decreased compared with control cells (Fig. 1D). This indicates that ACAA1 may have a substrate preference for medium-chain DCAs and would implicate SCPx in the metabolism of long-chain DCAs. Unexpectedly, DCA β-oxidation was normal in *SCP2* single KO cells (Fig. 1D). DCA β-oxidation was completely defective in *ACAA1/SCP2* DKO cells (Fig. 1D). These results show that ACAA1 and SCPx have overlapping functions in DCA metabolism, with a more important role of ACAA1 in the β-oxidation of medium-chain DCAs.

### EHHADH plays an important role in medium-chain DCA metabolism

The role of EHHADH in DCA β-oxidation is well documented (Ding et al. 2013; Dirkx et al. 2007; Ferdinandusse et al. 2004; Houten et al. 2012; Nguyen et al. 2008), but the data on the levels of DCAs in liver tissue and urine are not consistent. Ding et al reported an accumulation of DCAs (C6-to C16-DCA) in the liver of *Ehhadh* KO mice (Ding et al. 2013). In contrast, Houten et al reported decreased C6DC-carnitine in plasma, liver and urine as well as decreased urinary adipic acid (C6-DCA) excretion in *Ehhadh* KO mice (Houten et al. 2012). However, the urine organic acid analysis was hampered by a lack of sample volume (Houten et al. 2012). To further clarify the consequences of EHHADH deficiency on urinary excretion of DCAs, we now provide a systematic quantification of DCAs (C6-, C8-, C10- and C12-DCA as well as their 3-hydroxy intermediates) in the urine of *Ehhadh* KO mice. To increase the flux through the ω-oxidation and peroxisomal FAO pathways, we collected the samples after food withdrawal as well as from mice with pharmacological (L-AC) or genetic inhibition (*Acadl* KO) of mitochondrial FAO.

*Ehhadh^+/−^* animals were crossed with *Acadl^+/−^* animals (both on a pure C57BL/6 background) to generate double heterozygous animals. A cross of double heterozygous mice did not produce the expected number of progeny, with less *Ehhadh/Acadl* double KO (DKO) animals than expected (Table S2A). Similarly, a cross of *Ehhadh^-−/−^ Acadl^+/−^* × *Ehhadh^−/−^ Acadl^+/−^* mice had a less than expected number of DKO animals (Table S2B). Gestational loss is part of the phenotype of the *Acadl* KO mouse model (Berger & Wood 2004; Kurtz et al. 1998) but has not been observed in the *Ehhadh* KO mouse model (Qi et al. 1999). Using different breeding strategies, we ultimately obtained 6 DKO animals. These mice appeared healthy. Upon overnight food withdrawal, DKO animals were hypoglycemic, similarly to *Acadl* KO mice (Fig. 2A). Increases in the liver and heart relative weights were more pronounced in the DKO mice (Fig. 2A).

**Fig. 2.**
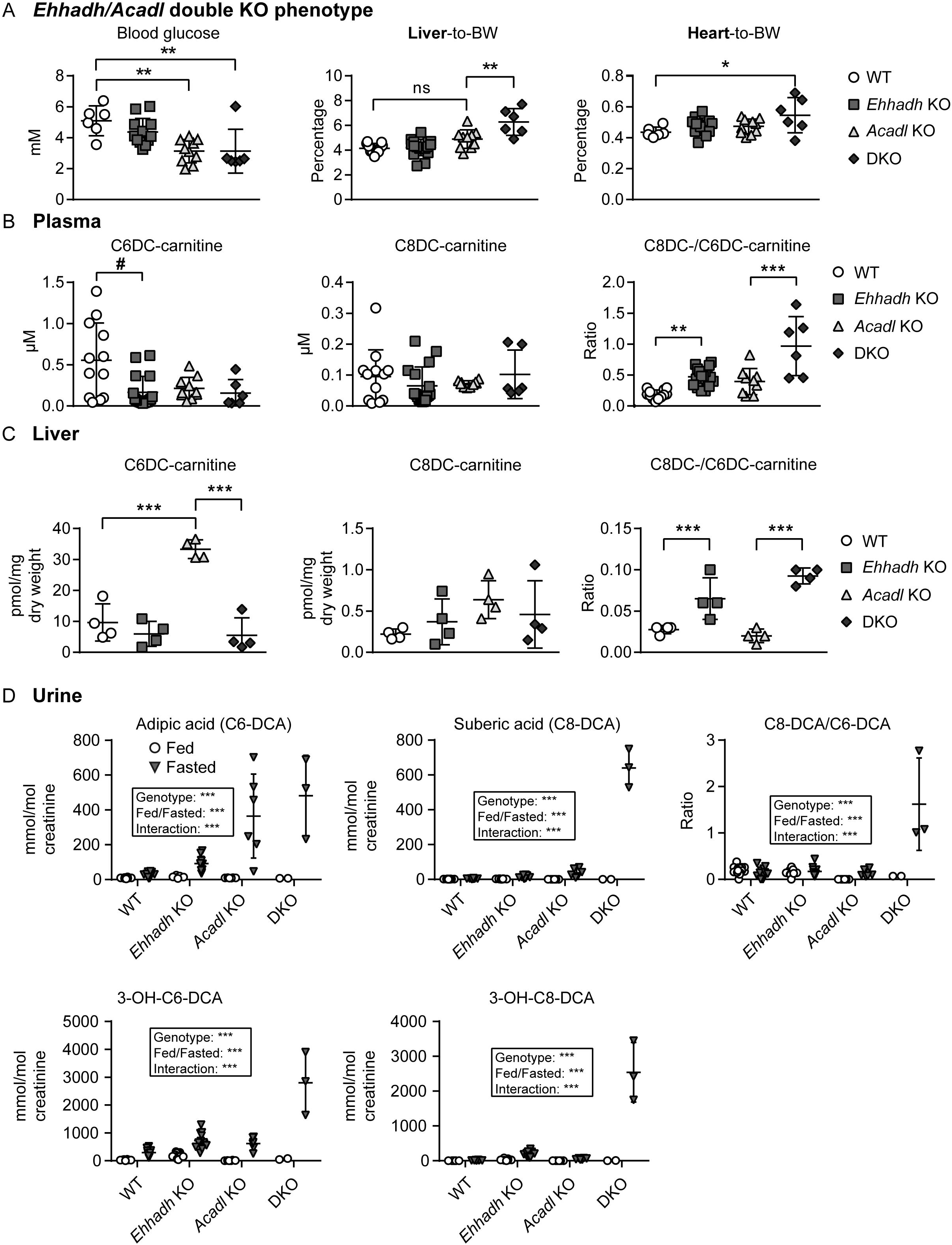
EHHADH plays an important role in medium-chain DCA metabolism. **A)** Blood glucose levels (mM), and relative liver-to-BW (body weight) and heart-to-BW percentages in WT (n=6-11), *Ehhadh* KO (n=12-17), *Acadl* KO (n=10) and *Ehhadh/Acadl* double KO mice (DKO, n=6). **B)** Plasma C6DC- and C8DC-carnitine (μM), and the C8DC/C6DC-carnitine ratio in WT (n=12), *Ehhadh* KO (n=17), *Acadl* KO (n=10) and DKO mice (n=6). **C)** Hepatic C6DC- and C8DC-carnitine (pmol/mg dry weight tissue), and the C8DC/C6DC-carnitine ratio in WT, *Ehhadh* KO, *Acadl* KO, and DKO mice, n=4 per genotype. **(D)** Urinary C6-DCA, C8-DCA, their 3-OH forms (mmol/mol creatinine), and the C8-DCA/C6-DCA ratio in WT (fed=20, fasted n=12), *Ehhadh* KO (fed=21, fasted n=12), *Acadl* KO (fed=6, fasted n=6) and DKO mice (fed=2, fasted n=3). Data are represented individually with the mean ± S.D. Statistical significance was tested using one-way ANOVA (**A-C**), two-way ANOVA with “Genotype” and “Fed/Fasted” as the two factors (**D**), or non-parametric Kruskal-Wallis test (C6DC- and C8DC-carnitine in **B**). After one-way ANOVA, Tukey’s multiple comparison test was performed *P < 0.05; **P < 0.01; ***P < 0.001. After non-parametric Kruskal-Wallis test, Dunn’s multiple comparison test was performed. # P < 0.05

We measured the acylcarnitine profile in the plasma (Table S3A) and liver (Table S3B) of these mice. A reproducible metabolite marker of *Ehhadh* KO and DKO mice was the increase in the ratio between C8DC- and C6DC-carnitine in plasma (Fig. 2B) and liver (Fig. 2C), which is mostly driven by decreased C6DC-carnitine levels. The ratio of these two metabolites can be viewed as a proxy for the β-oxidation of C8-DCA into C6-DCA. These data are consistent with the role of EHHADH in DCA metabolism and replicates our previous findings (Houten et al. 2012). *Acadl* KO mice present a characteristic acylcarnitine profile caused by the defect in mitochondrial FAO with a prominent elevation of C14:1-carnitine, accompanied by secondary carnitine deficiency (Bakermans et al. 2013; Kurtz et al. 1998; Ranea-Robles et al. 2020). However, no major differences were detected in the plasma or liver acylcarnitine profile of DKO mice compared with *Acadl* KO mice (Table S3A, S3B). Similar changes were observed in the acylcarnitine profile of *Ehhadh* KO mice after mitochondrial FAO inhibition with LAC (Violante et al. 2019). We observed an increase in C8DC-/C6DC-carnitine in the plasma and liver of L-AC-treated *Ehhadh* KO mice (Supplementary Fig. S2A, S2B). Overall, the consequences of a combined EHHADH and LCAD defect appear limited to DCA metabolism, underscoring the importance of EHHADH and peroxisomes in this metabolic pathway.

Medium-chain dicarboxylic aciduria is a prominent finding in the urine from fasted mice, especially in mice with a defect in mitochondrial FAO, including the *Acadl* KO mouse (Cox et al. 2001). We found elevated adipic acid (C6-DCA) levels in the urine of fasted *Ehhadh* KO, *Acadl* KO and DKO mice. The excretion of adipic acid was highest in *Acadl* KO and DKO mice (Fig. 2D). Although suberic acid (C8-DCA) was already increased in fasted *Acadl* KO mice, this increase was very notable in fasted DKO mice (Fig. 2D). Consequently, there was a significant increase in the ratio between urinary C8-DCA and C6-DCA in fasted DKO mice compared with single *Ehhadh* KO or *Acadl* KO mice (Fig. 2D). We also detected elevated levels of sebacic acid (C10-DCA) and dodecanedioic acid (C12-DCA) in the urine of fasted DKO mice (Supplementary Fig. S2C). The increase in adipic acid in the urine of *Ehhadh* KO mice is consistent with the reported elevation of this metabolite in liver extracts (Ding et al. 2013).

Even more remarkable changes were observed for the 3-hydroxy DCAs. 3-Hydroxyadipic acid (3-OH-C6-DCA) was elevated in the urine of fasted *Ehhadh* KO and *Acadl* KO mice, but most prominently in DKO mice (Fig. 2D). Although already increased in fasted *Ehhadh* KO mice, 3-hydroxysuberic (3-OH-C8-DCA) and 3-hydroxysebacic acid (3-OH-C10-DCA) levels were very high in DKO mice (Fig. 2D and Supplementary Fig. S2C). 3-Hydroxydodecanedioic acid (3-OH-C12-DCA) levels were generally low and not reliably detected. These findings were replicated in the urine of *Ehhadh* KO animals treated with L-AC (Supplementary Fig. S2D). From our data, we conclude that EHHADH plays an important role in medium-chain DCA metabolism signified by a prominent 3-hydroxydicarboxylic aciduria and an increased C8DC-to C6DC ratio.

### EHHADH deficiency reduces the β-oxidation of [U-^13^C]-C12-DCA in mouse liver slices

Ding et al. showed a defect in C12-DCA degradation and C6-DCA formation in liver homogenates from *Ehhadh* KO mice (Ding et al. 2013). We further investigated this defect using a precision-cut liver slice (PCLS) culture system. PCLS are viable *ex vivo* explants of mouse liver that are metabolically active and reduce the amount of substrate needed for metabolic tracing studies. PCLS also represent a more relevant physiological model compared with isolated cultured cells and liver homogenates (De Graaf et al. 2010; van de Bovenkamp et al. 2005). DCA metabolism was probed with [U-^13^C]-C12-DCA, which was added to the culture media of the PCLS from WT and *Ehhadh* KO mice. We then measured the amount of C6-DCA (M+6), C8-DCA (M+8), and their M+6 and M+8 carnitine conjugates, which reflect 3 or 2 completed cycles of DCA β-oxidation, respectively. We observed an increased ratio of [U-^13^C]-labeled C8-to C6-DCA in the media from *Ehhadh* KO liver slices (Fig. 3A). We also observed elevated levels of ^13^C-labeled 3-hydroxysuberic acid (3-OH-C8-DCA, M+8) in the media of *Ehhadh* KO liver slices compared with WT liver slices (Fig. 3B). The ratio of [U-^13^C]-labeled C8DC-to C6DC-carnitine tended to increase in the media of *Ehhadh* KO liver slices compared with those obtained from WT mice (Fig. 3C). Combined, these data further illustrate the important role of EHHADH in the hepatic β-oxidation of medium-chain DCAs.

**Fig. 3.**
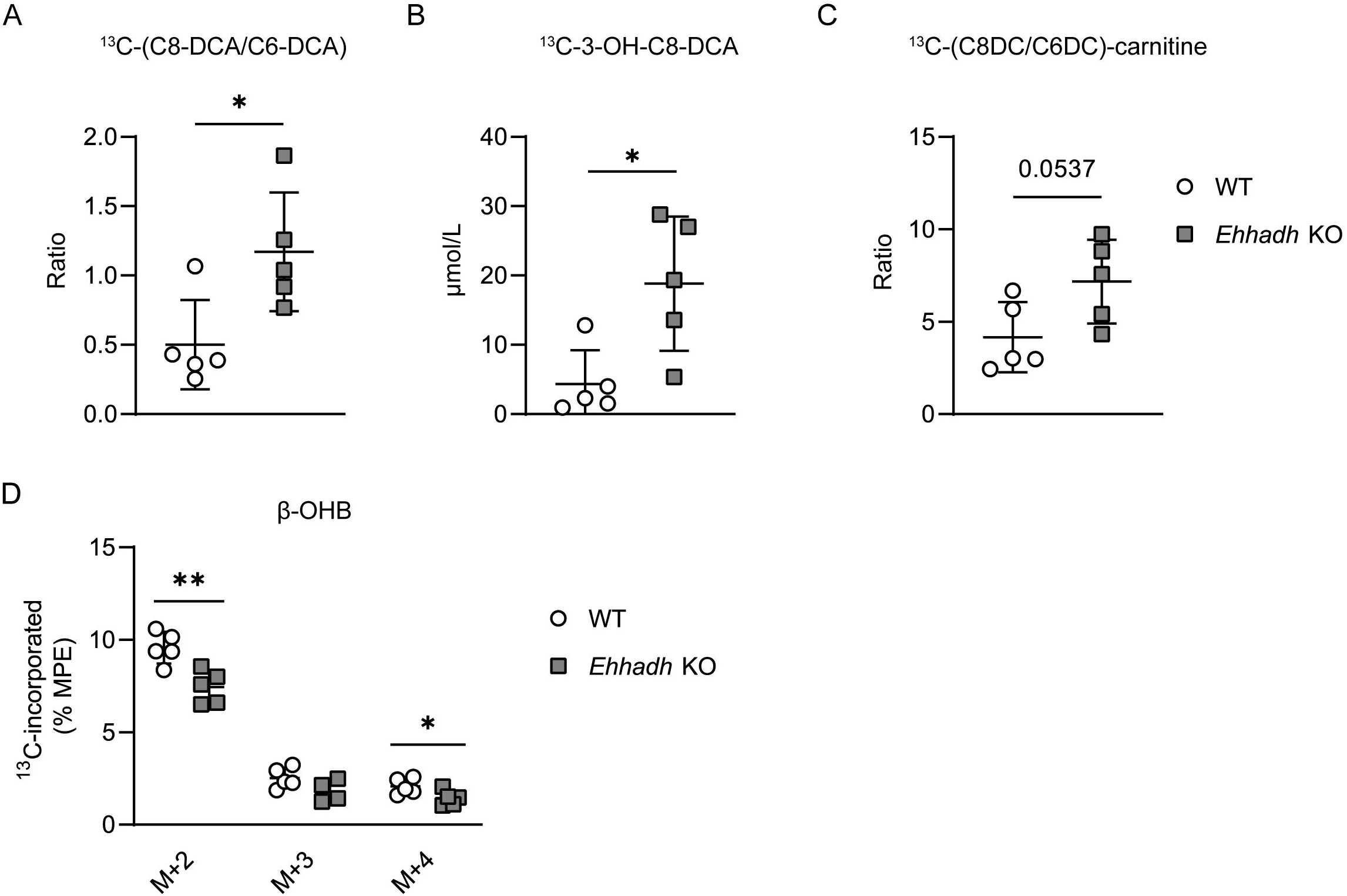
Metabolic tracing of [U-^13^C]-C12-DCA in precision-cut mouse liver slices. (**A-C**) Measured [U-^13^C]-labeled C6-DCA, C8-DCA, and 3-OH-C8-DCA (μmol/L) in media after 24-hr incubation of WT and *Ehhadh* KO liver slices with [U-^13^C]-C12-DCA, and calculation of the corresponding C8DCA-to C6-DCA (**A**) and C8-DC-to C6-DC-carnitine (**B**) ratios. **D**) MPE (molar percent enrichment) of individual ^13^C-labeled mass isotopomers of β-OHB in liver slices after 24-hr incubation with [U-^13^C]-C12-DCA. Data are represented individually with the mean ± S.D. Statistical significance was tested using unpaired t-test with Welch’s correction. *P < 0.05; **P < 0.01

To reveal the metabolic destination of the DCA carbons after peroxisomal β-oxidation, we measured the ^13^C-enrichment in TCA cycle intermediates, amino acids, and the ketone body β-hydroxybutyrate (β-OHB) in WT and *Ehhadh* KO liver slices after incubation with [U-^13^C]-C12-DCA (Zhang et al. 2015). We observed a ^13^C-enrichment in acetyl-CoA and TCA cycle intermediates, including succinate M+2 and M+4 (Supplementary Fig. S3). We noted a high enrichment in β-OHB M+2 (Fig. 3D) closely resembling the enrichment of acetyl-CoA. The ^13^C-enrichment in β-OHB (M+2 and M+4) was decreased in *Ehhadh* KO liver slices (Fig. 3D). These observations suggest that acetyl-CoA products derived from peroxisomal DCA β-oxidation can be shuttled to mitochondria for further metabolism such as oxidation or ketogenesis. The deficiency of EHHADH may decrease peroxisomal acetyl-CoA production from DCAs.

### Perturbation of hepatic cholesterol homeostasis in *Ehhadh* KO mice

To shed more light on the physiological functions of DCA metabolism, we performed RNA sequencing analysis of RNA isolated from the livers of WT and *Ehhadh* KO animals after an overnight food withdrawal. We only found 17 differentially expressed genes (DEG) by DESeq2 (12 up and 5 down), of which 2 were also detected by limma technology (Fig. 4A, Table S4A and S4B). The low number of DEG is consistent with the absence of detectable phenotypic defects in the *Ehhadh* KO mouse (Qi et al. 1999). Interestingly, within the DEG, we noted two genes involved in cholesterol biosynthesis; *Hmgcr* (hydroxymethylglutaryl-CoA reductase) and *Lss* (lanosterol synthase) (Fig. 4A). Next, we used Gene Set Enrichment Analysis (GSEA), which revealed a significant upregulation of genes involved in cholesterol homeostasis (Fig. 4B and Table S4C). No other pathways were significantly enriched (Table S4C).

**Fig. 4.**
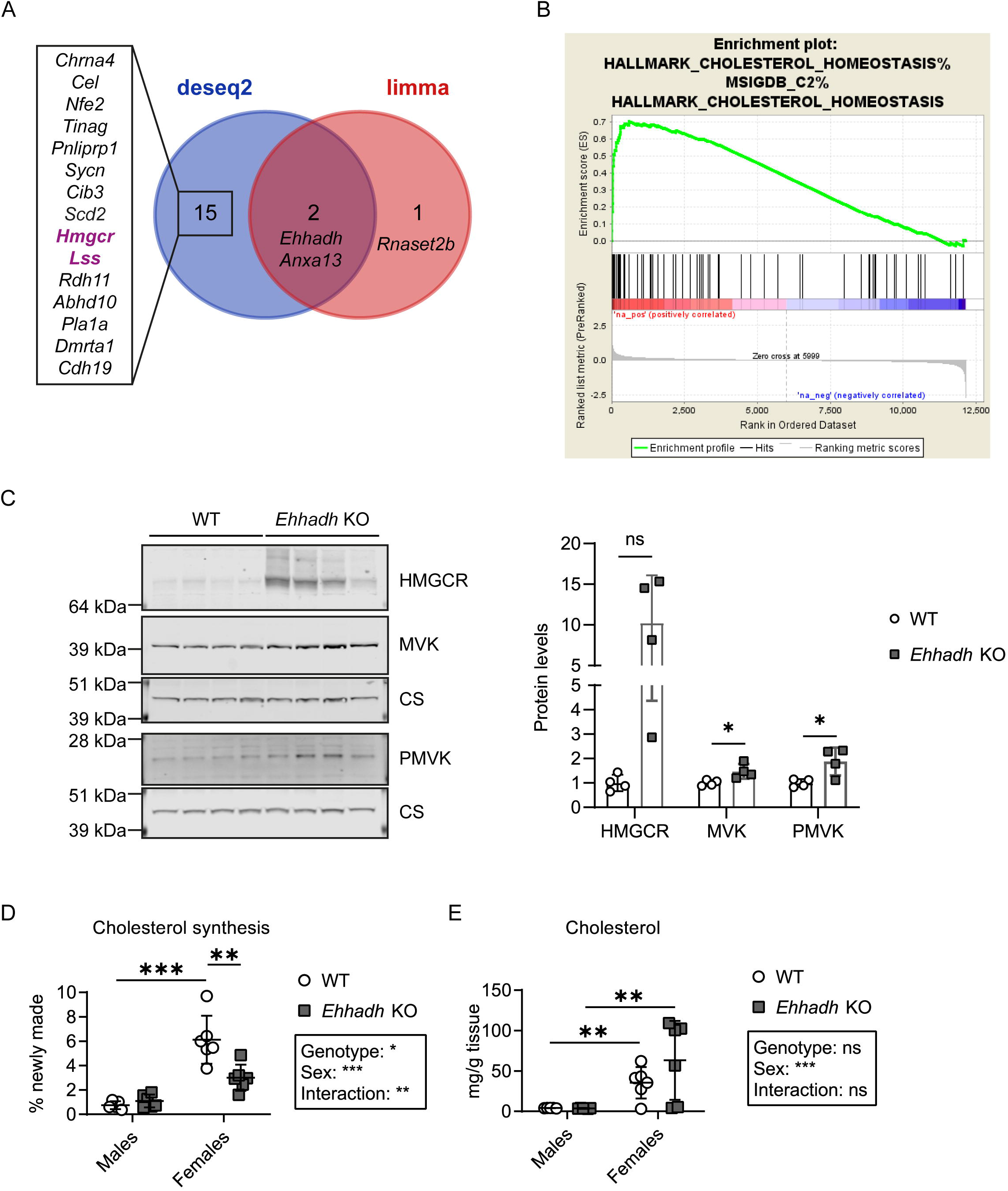
Perturbation of hepatic cholesterol homeostasis in *Ehhadh* KO mice. **A)** Venn diagram comparing the number of DEG detected by both DESeq2 and limma technologies. All gene names are provided, and those belonging to the cholesterol biosynthesis pathway are highlighted. **B)** Enrichment plot for the HALLMARK CHOLESTEROL HOMEOSTASIS gene set in the molecular signature database (M5892). The graph shows the profile of the running Enrichment Score (ES) and the positions of gene set members on the fold-change rank-ordered list. Overall the NES was 2.338538 with an FDR q-value of 0. **C)** Immunoblots for HMGCR, MVK, PMVK and the loading control citrate synthase (CS) in WT and *Ehhadh* KO mice (n=4 per genotype), and the corresponding quantification. **D)** Cholesterol synthesis was calculated from the incorporation of deuterium isotopes (^2^H) into cholesterol in the liver. **E)** Hepatic cholesterol content (mg/g tissue) was measured by mass spectrometry. Data are represented individually with the mean ± SD. Statistical significance was tested using unpaired t-test with Welch’s correction (**C**) or two-way ANOVA with “Genotype” and “Sex” as the two factors followed by Tukey’s multiple comparisons test (**D, E**). *P < 0.05; **P < 0.01; ***P < 0.001.

Given the small fold change of the upregulation of genes involved in cholesterol homeostasis, we studied protein expression. We found that HMGCR, which catalyzes the rate-limiting step in cholesterol biosynthesis, was increased 10.2-fold in *Ehhadh* KO livers (Fig. 4B). The increase in HMGCR protein was much larger than the increase in *Hmgcr* mRNA, which may be explained by well-documented posttranscriptional regulatory mechanisms of HMGCR (Goldstein & Brown 1990; Jun et al. 2020). Mevalonate kinase (MVK, 1.5-fold) and phosphomevalonate kinase (PMVK, 1.9-fold) were also increased in *Ehhadh* KO livers (Fig. 4B), but the fold-change was much smaller when compared to HMGCR. The elevation of HMGCR, MVK and PMVK in *Ehhadh* KO mice was reproduced in an independent set of WT and *Ehhadh* KO mice (Supplementary Fig. S4A). HMGCR protein levels were increased both in male and female *Ehhadh* KO mice (Supplementary Fig. S4B), as well as in *Ehhadh* KO animals treated with L-AC (Supplementary Fig. S4C). These results show that cholesterol homeostasis is perturbed in *Ehhadh* KO mice.

To support the role of peroxisomal metabolism in cholesterol homeostasis, we measured *de novo* lipogenesis (DNL) and cholesterol biosynthesis in WT and *Ehhadh* KO mice using the ^2^H_2_O method (Bederman et al. 2009; Brunengraber et al. 2003). We found that the amount of hepatic cholesterol and the rate of hepatic cholesterol synthesis were sexually dimorphic, with higher rates and increased levels of cholesterol in female mice (Fig. 4C, D). Cholesterol synthesis was decreased in female *Ehhadh* KO and not changed in male *Ehhadh* KO mice compared to sex-matched controls (Fig. 4C). We did not find any changes in DNL in *Ehhadh* KO mice compared with WT mice (Supplementary Fig. S4D, S4E). Thus, the apparent increase in the mRNA and protein expression of enzymes involved in cholesterol synthesis does not translate into increased cholesterol synthesis. In fact, cholesterol synthesis was decreased in female *Ehhadh* KO mice. These data further indicate a perturbation of cholesterol homeostasis in *Ehhadh* KO mice.

## Discussion

In this study, we provide a detailed characterization of the contribution of individual peroxisomal transporters and enzymes to the β-oxidation of DCAs in a human HEK-293 cell line model. We also provide further evidence supporting the specific role of EHHADH in the hepatic metabolism of DCAs, and its functional relevance as a modulator of hepatic cholesterol metabolism.

Our study provides more definite genetic proof of the essential role of peroxisomes and the peroxisomal enzyme EHHADH in DCA β-oxidation, supporting previous research that also pointed to that direction (Ding et al. 2013; Dirkx et al. 2007; Houten et al. 2012; Nguyen et al. 2008; Poosch & Yamazaki 1989; Suzuki et al. 1989). Based on our experiment in HEK-293 cells, we propose that the peroxisomal pathway of DCA β-oxidation is formed by the transporter ABCD3, the acyl-CoA oxidase ACOX1, the bifunctional enzyme HSD17B4 (with a small contribution of EHHADH due to low expression), and the thiolases ACAA1 and SCPx. Overexpression of EHHADH can functionally compensate for HSD17B4 deficiency, indicating that both EHHADH and HSD17B4 are able to handle DCAs and that their functional role in this process may be mainly driven by tissue expression levels. Our results also provide evidence for a partially overlapping function of ACAA1 and SCPx in DCA β-oxidation, with ACAA1 being more important in the β-oxidation of medium-chain DCAs. We speculate that additional players involved in DCA β-oxidation are a dicarboxylyl-CoA synthetase needed for the activation of DCAs into DCA-CoAs (ACSL1, acyl-CoA synthetase long chain family member 1) and subsequent uptake by peroxisomes as well as CROT (peroxisomal carnitine O-octanoyltransferase) and ACOT4 (Houten et al. 2012, 2020; Hunt et al. 2006; Vamecq et al. 1985; van Roermund et al. 2020; Westin et al. 2005). The role of these enzymes in DCA metabolism should be addressed in future studies. Finally, even though our study in HEK-293 cells did not find any role for mitochondrial FAO in DCA β-oxidation, we cannot exclude that under pathophysiological conditions such as an impairment of peroxisomal FAO, DCAs are metabolized in the mitochondria. This hypothesis deserves further investigation using models of peroxisomal dysfunction.

Remarkably, no human EHHADH defect has been described to date and, therefore, the *Ehhadh* KO mouse remains the best tool to study EHHADH’s role in mammalian physiology. We provide additional evidence for the role of EHHADH in DCA β-oxidation by the identification of medium-chain 3-hydroxydicarboxylic aciduria and an increased ratio of C8-DCA to C6-DCA and C8DC-to C6DC-carnitine in *Ehhadh* KO mice. We speculate that a similar metabolite pattern could indicate a human deficiency in a case with potentially pathogenic biallelic *EHHADH* variants. However, 3-hydroxydicarboxylic aciduria by itself is not a specific marker as it frequently accompanies dicarboxylic aciduria, for example, during fasting (Tserng & Jin 1991b; Tserng et al. 1989). In addition, mitochondrial long-chain 3-hydroxyacyl-CoA dehydrogenase (LCHAD) deficiency, as well as other conditions, cause 3-hydroxydicarboxylic aciduria (Bennett & Sherwood 1993; Bennett et al. 1994; Bergoffen et al. 1993; Mayatepek et al. 1995; Tserng et al. 1991). Ehhadh KO mice have no obvious hepatic disease phenotype under normal conditions, but feeding these animals with a diet rich in medium-chain fatty acids led to liver inflammation, fibrosis, and death (Ding et al. 2013). We speculate that a potential clinical presentation of human EHHADH deficiency may be hepatic toxicity of medium-chain fat rich foods (e.g. coconut oil) or supplements such as medium-chain triglyceride oil.

Our results also indicate that DCA β-oxidation is not completely defective in *Ehhadh* KO mice but appears to have a limited capacity for medium-chain DCAs. There are several lines of evidence to support this. First, our *in vitro* experiments have clearly demonstrated that EHHADH and HSD17B4 display functional redundancy in the metabolism in DCAs, and their contribution to this process appears to depend on their expression level. This further highlights the role of EHHADH in DCA β-oxidation, given its high expression levels in liver and kidney. Second, and somewhat counterintuitively, *Ehhadh* KO mice are able to generate medium-chain DCAs, including adipic acid, which is considered an end product of peroxisomal DCA β-oxidation. However, the fact that the highest levels of intermediates are observed for suberic acid and 3-hydroxysuberic acid, which are both not typically viewed as final products of DCA metabolism, suggests that the pathway is saturated. Indeed, the ratio of C8DC over C6DC is consistently higher in *Ehhadh* KO animals suggesting that the main bottleneck is at the level of C8DC β-oxidation. Lastly, stable isotope tracing in liver slices using ^13^C-C12-DCA as a precursor reproduced the increased ratio of C8-DCA over C6-DCA for the ^13^C labeled products in the *Ehhadh* KO media compared to WT media. Ultimately our observations are consistent with a model where all DCA β-oxidation intermediates compete for a common set of enzymes. Such competition becomes problematic at saturating conditions (van Eunen et al. 2013), which may be because EHHADH is deficient and HSD17B4 is the only available enzyme and/or because DCA metabolism is induced due to fasting or a defect in mitochondrial FAO.

Remarkably, liver and plasma C6DC-carnitine levels are generally decreased in *Ehhadh* KO mice, whereas urine and liver adipic acid levels are increased. This suggests that adipoyl-CoA metabolism differs between *Ehhadh* KO and WT mice. Specifically, conversion to C6DC-carnitine by CROT is decreased, whereas thiolytic cleavage is increased in *Ehhadh* KO mice. Given that *Ehhadh* KO mice do not display secondary carnitine deficiency, we speculate that increased thiolytic cleavage is the likely culprit. Indeed, it is known that ACOT8, which displays activity with medium-chain DCAs, is inhibited by free CoA (Hunt et al. 2002; Westin et al. 2005). We suggest that an accumulation of DCA β-oxidation intermediates in *Ehhadh* KO mice due to saturation of the β-oxidation pathway may lead to reduced availability of peroxisomal free CoA (i.e. CoA trapping), which would stimulate ACOT8 activity and as such generate higher levels of free DCA intermediates.

The finding that 3-hydroxyadipic acid accumulates in fasted *Ehhadh* KO, *Acadl* KO and *Acadl/Ehhadh* double KO mice is notable because it indicates that C6-DCA can undergo another round of peroxisomal β-oxidation, which would involve EHHADH and yield the anaplerotic substrate succinate. The low but detectable labeling of TCA cycle intermediates after incubating WT and *Ehhadh* KO liver slices with [U-^13^C]-C12-DCA further supports this possibility. The capacity of peroxisomes to generate succinate is also supported by the work on ACOT4, a peroxisomal acyl-CoA thioesterase that displays optimal activity with succinyl-CoA (Hunt et al. 2006; Westin et al. 2005). Even though some studies using ^13^C-labeled DCAs were not able to detect metabolism beyond adipic acid (Cerdan et al. 1988), other studies clearly established anaplerosis from C12-DCA (Jin & Tserng 1991; Jin et al. 2015; Tserng & Jin 1991a). The anaplerotic potential of peroxisomal DCA β-oxidation products in different physiological situations remains to be established. We provide evidence in this study of the feasibility of mouse liver slice culture to perform isotope tracer studies to study metabolic reactions, which could be used to investigate this aspect of peroxisomal metabolism.

Finally, our study shows that EHHADH deficiency perturbs cholesterol biosynthesis. We observed an induction of cholesterol biosynthesis enzymes at the mRNA and protein level, including a pronounced increase of HMGCR, the rate-limiting enzyme in cholesterol biosynthesis. An increase in the expression of cholesterol biosynthetic genes is usually mediated by decreased levels of cholesterol and non-sterol isoprenoids leading to the activation of pathways, such as the proteolytic cleavage of membrane-bound sterol regulatory element binding protein 2 (SREBP2), which aim to restore cholesterol levels (Brown & Goldstein 1997). Despite this induction of cholesterol biosynthesis genes, cholesterol biosynthesis, as measured by the ^2^H_2_O method, was reduced in female and not changed in male *Ehhadh* KO mice. Combined, these data indicate that cholesterol biosynthesis is decreased due to EHHADH deficiency leading to a compensatory increase in cholesterol biosynthetic capacity via the SREBP2 pathway.

We propose that peroxisomal β-oxidation is an important source of cytosolic acetyl-CoA that can be used for cholesterol and fatty acid biosynthesis. Thus, a defect in peroxisomal β-oxidation limits the availability of this biosynthetic precursor. Peroxisomal acetyl-CoA destined for use in the cytosol is presumably released in the form of acetate (Kasumov et al. 2005; Leighton et al. 1989), which is supported by the presence of peroxisomal thioesterases with an affinity for short chain acyl-CoA molecules such as ACOT8 and ACOT12 (Hunt et al. 2014). Previous studies have shown that administration of DCAs leads to cytosolic acetyl-CoA production (Jin et al. 2015; Leighton et al. 1989), which can be used for the biosynthesis of cholesterol (Dietschy & McGarry 1974; Kondrup & Lazarow 1985). We assessed the fate of acetyl-CoA produced from [U-^13^C]-C12-DCA in mouse liver slices from WT and *Ehhadh* KO mice. Although the acetyl-CoA pool was enriched, we were not able to detect ^13^C-enrichment in cholesterol perhaps due to a low overall cholesterol synthesis rate in the mouse liver slices. We did, however, obtain evidence that peroxisomal DCA β-oxidation provides acetyl-CoA units for other biochemical pathways such as ketogenesis. The reduced ^13^C-enrichment observed in the ketone body β-OHB in *Ehhadh* KO mouse liver slices compared to WT slices after adding the ^13^C-C12-DCA tracer supports the important role of EHHADH in DCA β-oxidation. Even though our results demonstrate that peroxisomal DCA β-oxidation yields acetyl-CoA that can be used for various metabolic processes, further experiments in relevant models of peroxisomal FAO deficiencies are warranted to assess this contribution under different physiologic conditions.

The role of peroxisomes in cholesterol metabolism remains a debated issue. *In vitro* and *in vivo* studies done in different models of peroxisomal deficiency have reported disturbances of cholesterol homeostasis and activation of the SREBP2 pathway (Charles et al. 2020; Fidaleo et al. 2011; Kovacs et al. 2004, 2009; Martens et al. 2008; Nicolas-Francès et al. 2014), in line with our findings in the liver of *Ehhadh* KO mice. Our observations are consistent with previous reports of reduced cholesterol synthesis rates in skin fibroblasts from patients with peroxisomal disorders (Hodge et al. 1991). In contrast, other reports have shown increased rates of cholesterol synthesis in *Pex2* KO and *Hsd17b4* KO mice (Kovacs et al. 2004; Martens et al. 2008). In these studies, cholesterol synthesis was increased when measured using labeled acetate. This increase, however, could be artifactual if the size of the acetate pool differs between WT mice and mice with a peroxisomal disorder. A decreased acetate pool in mice with peroxisomal disorders would lead to a higher enrichment with the same dose of labeled acetate, and a perceived increase in cholesterol synthesis. The ^2^H_2_O method that we employed does not have this potential drawback and directly implicates peroxisomal β-oxidation in the regulation of cholesterol homeostasis.

In summary, our results show that EHHADH has an important role in hepatic DCA metabolism and highlight the role of peroxisomal FAO in the regulation of cholesterol biosynthesis. Further studies are necessary to understand the physiological significance of these findings.

## Acknowledgements

We thank Dr. Hans Waterham for providing the MVK and PMVK antibodies, Jacob Hagen for the RNAseq analysis, the Flow Cytometry CoRE at the Icahn School of Medicine at Mount Sinai for their technical assistance with cell sorting, and Purvika Patel and Dr. Brandon Stauffer (Sema4) for their kind and excellent technical assistance with the clinical biochemical analyses.

Research reported in this publication was supported by the National Institute of Diabetes and Digestive and Kidney Diseases of the National Institutes of Health under Award Number R01DK113172. The content is solely the responsibility of the authors and does not necessarily represent the official views of the National Institutes of Health.

