## Supplementary figures and images for "Murine deficiency of peroxisomal L-bifunctional protein (EHHADH) causes medium-chain 3-hydroxydicarboxylic aciduria and perturbs hepatic cholesterol homeostasis"

### Figure S1

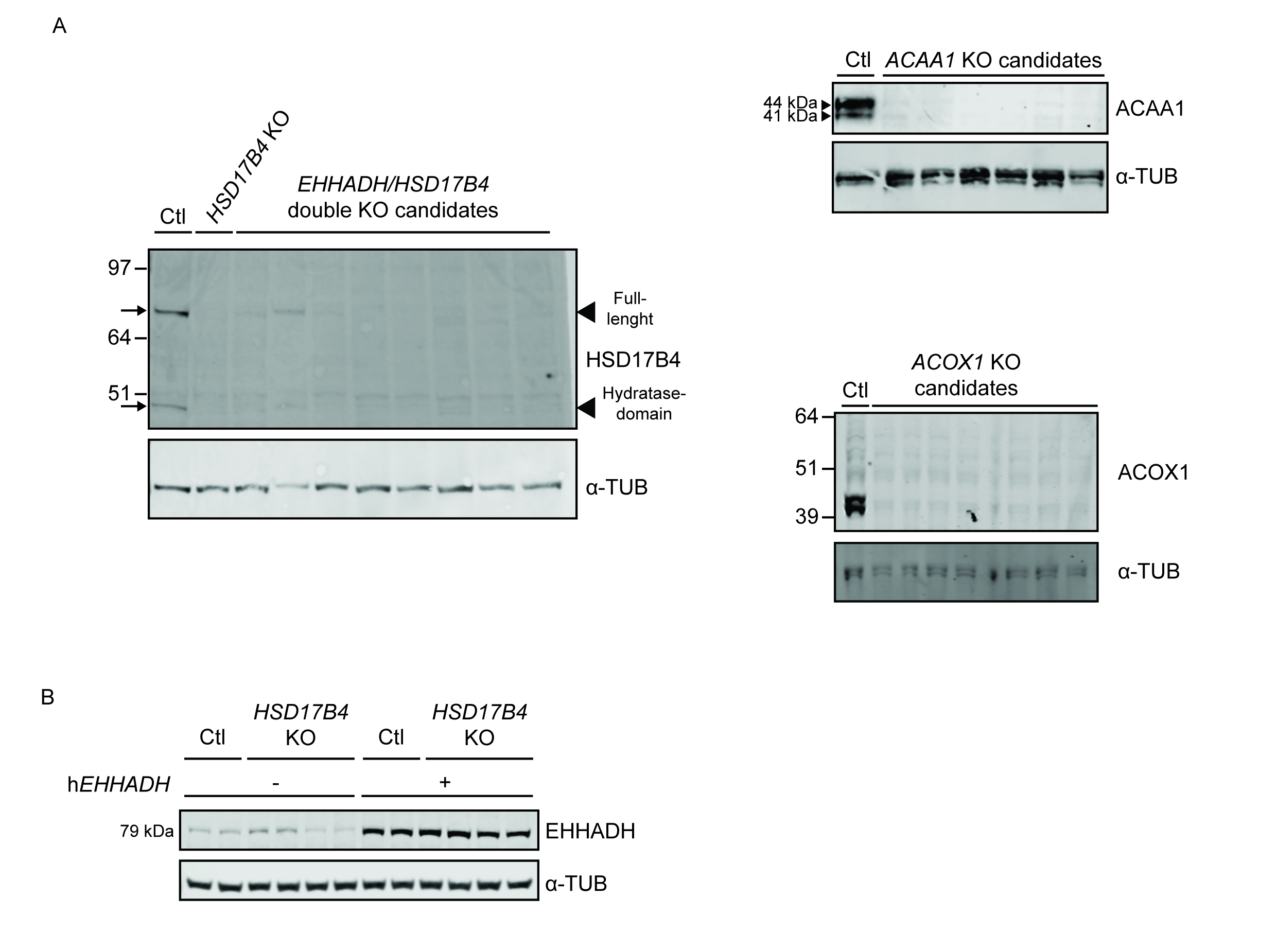

### Figure S2

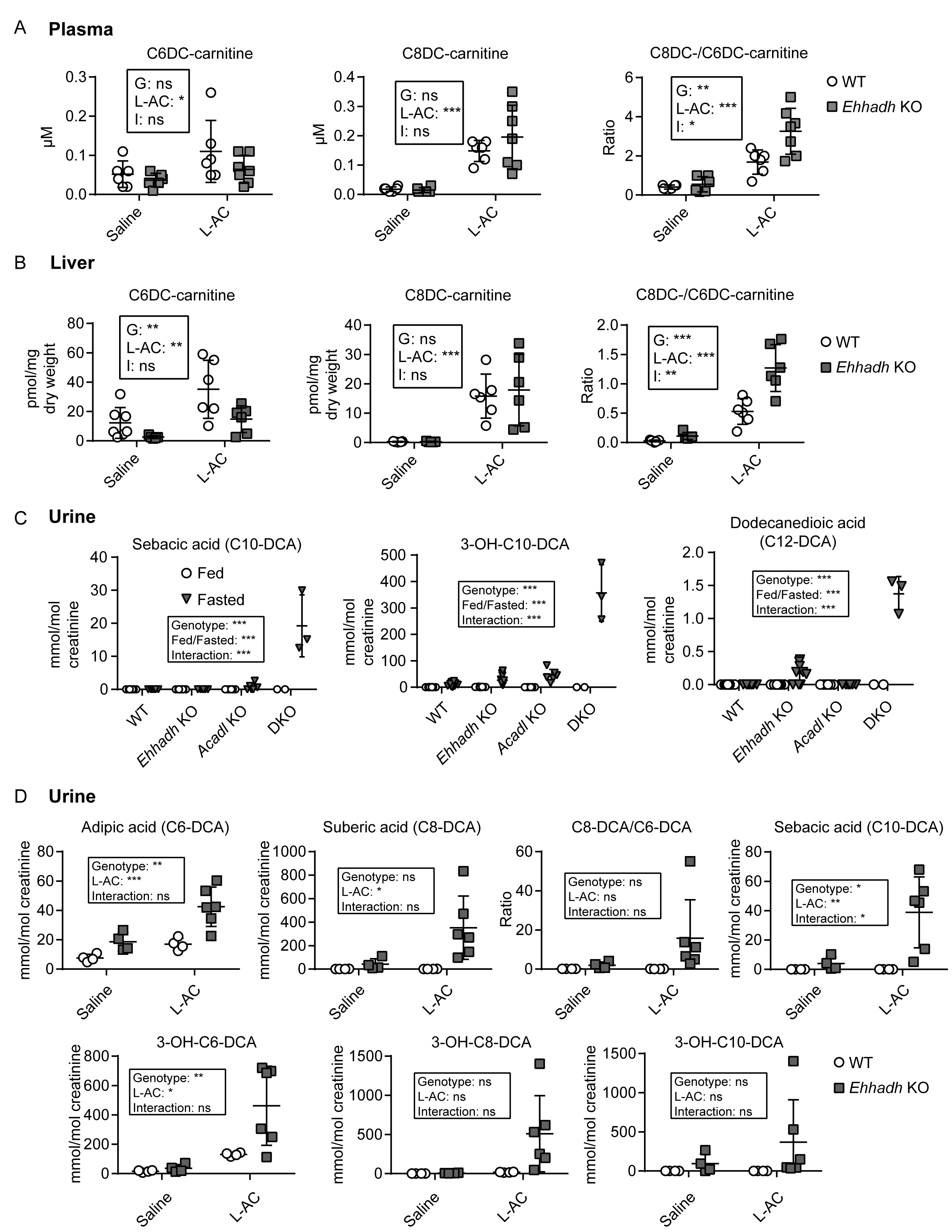

### Figure S3

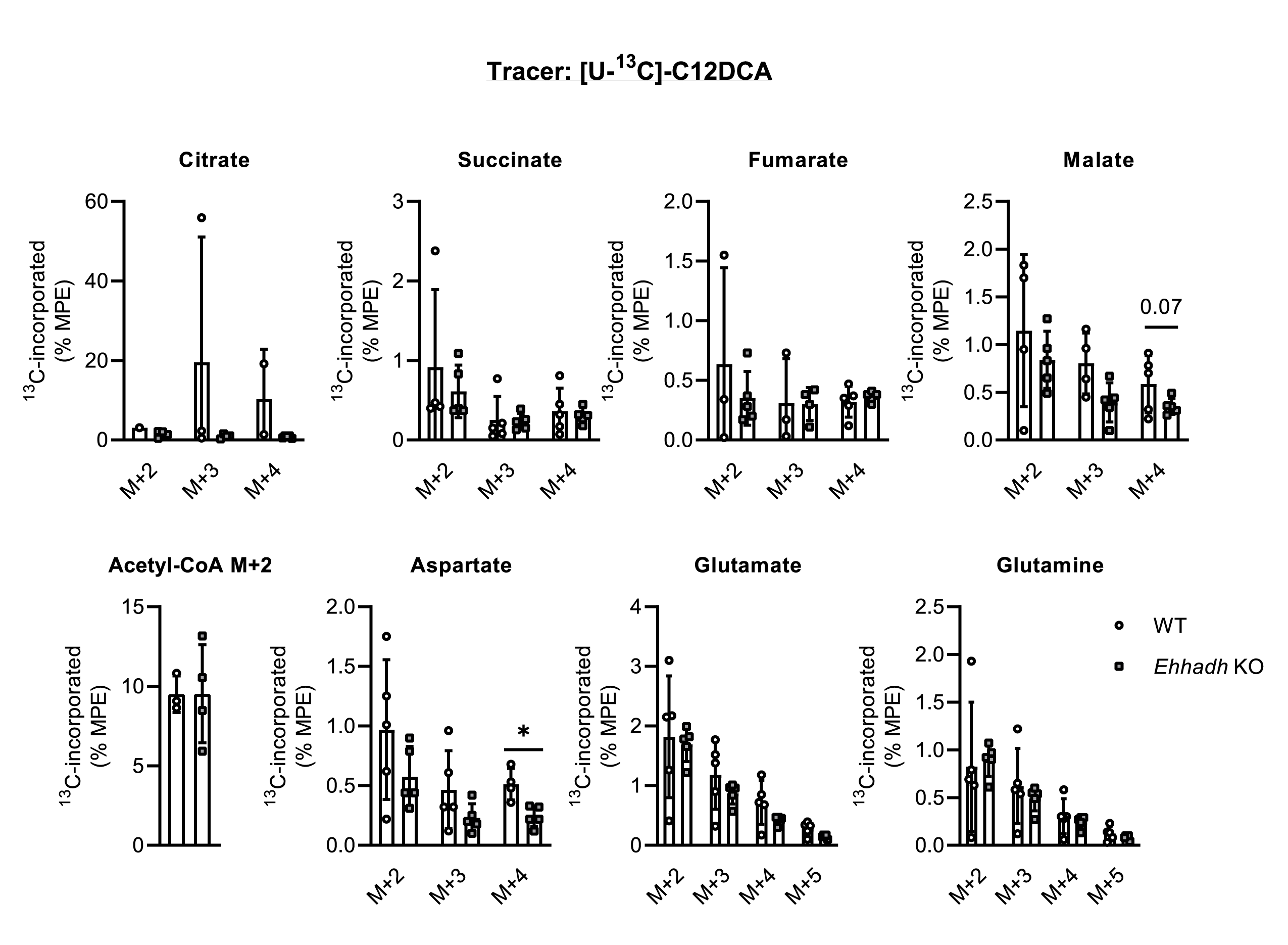

### Figure S4

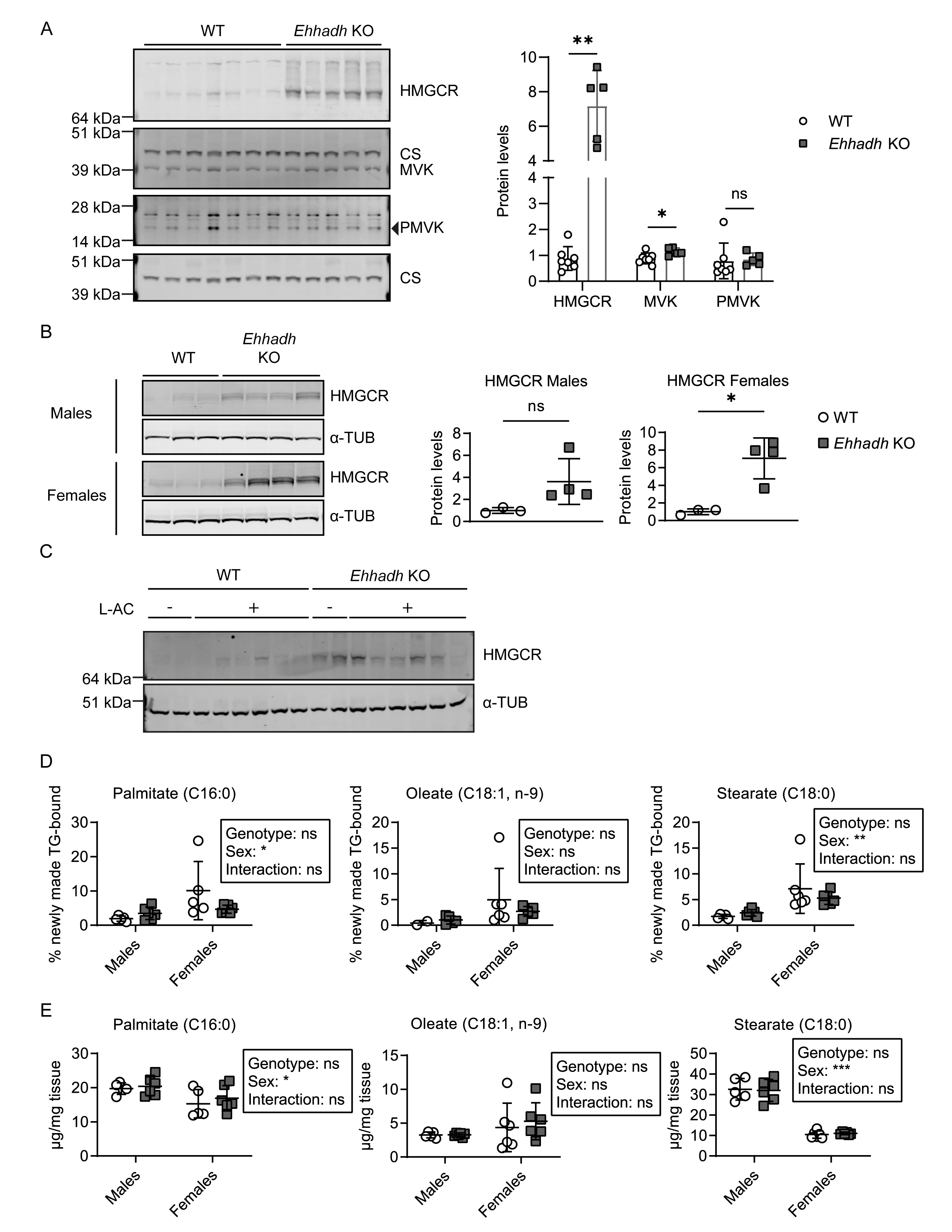
