## Supplementary information for "Murine deficiency of peroxisomal L-bifunctional protein (EHHADH) causes medium-chain 3-hydroxydicarboxylic aciduria and perturbs hepatic cholesterol homeostasis"

**Supplementary figure legends**

**Supplementary Fig. S1**. **Generation and validation of HEK-293 cell lines with peroxisomal enzyme deficiencies.**

**A)** HSD17B4, ACAA1 and ACOX1 immunoblots to validate CRISPR-Cas9 genome editing in HEK-293 cells. **B**) EHHADH immunoblot comparing control (Ctl) and *HSD17B4* KO cells with (+) or without (-) overexpression of EHHADH. Alpha-tubulin (α-TUB) was used as a loading control.

**Supplementary Fig. S2.** **EHHADH plays an important role in medium-chain DCA metabolism.**

**A)** Plasma C6DC- and C8DC-carnitine (µM), and the C8DC/C6DC-carnitine ratio in WT and *Ehhadh* KO mice with or without L-aminocarnitine (L-AC). n=6-7 per group. **B)** Hepatic C6DC- and C8DC-carnitine (pmol/mg dry weight tissue), and the C8DC/C6DC-carnitine ratio in WT and *Ehhadh* KO mice with or without L-AC (n=5-7 per group). **(C)** Urinary C10-DCA, 3-OH-C10-DCA, and C12-DCA (mmol/mol creatinine) in WT (fed=20, fasted n=12), *Ehhadh* KO (fed=21, fasted n=12), *Acadl* KO (fed=6, fasted n=6) and *Ehhadh*/*Acadl* double KO mice (DKO, fed=2, fasted n=3). **(D)** Urinary C6-DCA and C8-DCA, their 3-OH forms (mmol/mol creatinine), and the C8-DCA/C6-DCA ratio in WT and *Ehhadh* KO mice with or without L-AC (n=4-6 per group). Data are represented individually with the mean ± S.D. Statistical significance was tested using two-way ANOVA with the corresponding factors as indicated on each graph. *P < 0.05; **P < 0.01; ***P < 0.001.

**Supplementary Fig. S3. Metabolic tracing of [U-^13^C]-C12-DCA in precision cut mouse liver slices.**

Molar percent enrichment (MPE) of individual ^13^C-labeled mass isotopomers of selected metabolites in liver slices after 24-hr incubation with [U-^13^C]-C12-DCA (n=5 per genotype). Samples with undetectable ^13^C-enrichment are not plotted. Data are represented individually with the mean ± S.D. Statistical significance was tested using unpaired t test with Welch’s correction. *P < 0.05

**Supplementary Fig. S4**. **Perturbation of hepatic cholesterol homeostasis in *Ehhadh* KO mice.**

**A**) Immunoblots of HMGCR, MVK, PVMK and the loading control citrate synthase (CS) in WT (n=7) and *Ehhadh* KO (n=5) mice, and the corresponding quantification. **B**) Immunoblots of HMGCR and the loading control alpha-tubulin (α-TUB) in male and female WT (n=3 per sex) and *Ehhadh* KO (n=4 per sex) mice, and the corresponding quantification. **C**) Immunoblots of HMGCR and the loading control alpha-tubulin (α-TUB) in WT and *Ehhadh* KO mice with or without L-AC (n=2 WT-Veh and *Ehhadh* KO –Veh, n=6 WT+L-AC and *Ehhadh* KO+L-AC), and the corresponding quantification. **D**) *De novo* lipogenesis was calculated from the incorporation of deuterium isotopes (^2^H) into the corresponding TG-bound fatty acids (palmitate, oleate, and stearate) in the liver. **E**) TG-bound fatty acids (palmitate, oleate, and stearate) content (µg/mg tissue) was measured by mass spectrometry. Data are represented individually with the mean ± SD. Statistical significance was tested using unpaired t test with Welch’s correction (**A, B**), or two-way ANOVA with genotype and sex as the two factors (**D, E**) . *P < 0.05; **P < 0.01; *** P < 0.001.

**Supplementary Tables**

Table S1. List of CRISPR-Cas9 KO cell lines generated with their guide sequences

Table S2. Genotype distributions obtained for breeding of *Ehhadh* / *Acadl* DKO animals. (A) Genotype distribution in a cross of *Ehhadh*^+/-^*Acadl*^+/-^ x *Ehhadh*^+/-^*Acadl*^+/-^ mice. (B) Genotype distribution in a cross of *Ehhadh*^-/-^*Acadl*^+/-^ x *Ehhadh*^-/-^*Acadl*^+/-^ mice. Supplied as excel tables.

Table S3. (A) Plasma acylcarnitine profile in WT, *Ehhadh*^-/-^, *Acadl*^-/-^, and DKO mice. (B) Hepatic acylcarnitine profile in WT, *Ehhadh*^-/-^, *Acadl*^-/-^, and DKO mice. Supplied as excel tables.

Table S4. RNAseq analysis of WT and *Ehhadh*^-/-^ liver. (A) Differentially expressed genes analysis by DESeq2. (B) Differentially expressed genes analysis by limma. (C) GSEA analysis. Supplied as excel tables.

| **Table S1.** List of newly generated KO cell lines using CRISPR-Cas9 genome editing. The guide sequences used are indicated. | | | |
| --- | --- | --- | --- |
| **Cell line** | **Gene(s)** | **Guide 1** | **Guide 2** |
| *ACOX1* KO | *ACOX1* | CCATCCGATACAGCGCTGTG | CATGTCGGATGGCTTGTGGT |
| *ACAA1* KO | *ACAA1* | TCTCTCGGCAGTCATGACCG | ATTCACGTCCTTGAGAACCG |
| *SCP2* KO and *ACAA1/SCP2* DKO | **SCP2* | ATTCAGGTGACTCTACCTGT | ATCCATACTTATTGATCAGG |
| *EHHADH/HSD17B4* DKO | **HSD17B4* | ATTGGGCCGAGCCTATGCCC | TGGCTTTTGCAGAAAGAGGA |
| **SCP2* was targeted in an *ACAA1* KO background to create *ACAA1/SCP2* DKO  ** *HSD17B4* was targeted in an *EHHADH* KO background to create *EHHADH/HSD17B4* DKO | | | |
